# Hawkmoths use wingstroke-to-wingstroke frequency modulation for aerial recovery to vortex ring perturbations

**DOI:** 10.1101/2020.12.07.413781

**Authors:** Jeff Gau, Ryan Gemilere, LDS-VIP (FM subteam), James Lynch, Nick Gravish, Simon Sponberg

## Abstract

Centimeter-scale fliers that combine wings with springy elements must contend with the high power requirements and mechanical constraints of flapping wing flight. Insects utilize elastic energy exchange to reduce the inertial costs of flapping wing flight and potentially match wingbeat frequencies to a mechanical resonance. Flying at resonance may be energetically favorable under steady conditions, but it is difficult to modulate the frequency of a resonant system. Evidence suggests that insects utilize frequency modulation over long time scales to adjust aerodynamic forces, but it remains an open question the extent to which insects can modulate frequency on the wingstroke-to-wingstroke timescale. If wingbeat frequencies deviate from resonance, the musculature must work against the elastic flight system, thereby potentially increasing energetic costs. To assess how insects address the simultaneous needs for power and control, we tested the capacity for wingstroke-to-wingstroke wingbeat frequency modulation by perturbing free hovering *Manduca sexta* with vortex rings while recording high-speed video at 2000 fps. Because hawkmoth flight muscles are synchronous, there is at least the potential for the nervous system to modulate frequency on each wingstroke. We observed *±* 16% wingbeat frequency modulation in just a few wing strokes. Via instantaneous phase analysis of wing kinematics, we found that over 85% of perturbation responses required active changes in motor input frequency. Unlike their robotic counterparts that explicitly abdicate frequency modulation in favor of energy efficiency, we find that wingstroke-to-wingstroke frequency modulation is an underappreciated control strategies that complements other strategies for maneuverability and stability in insect flight.

## 1 Introduction

From undulatory swimming to legged locomotion, animals must contend with the dual challenges of control and energetics while constrained by the physics of their mechanical bodies [1, 2]. These challenges are particularly acute for flying animals, which must simultaneously generate sufficient lift and counteract inherent instabilities to remain airborne. Like resonant oscillators, insects store excess kinetic energy during a wing stroke in spring-like structures and return this energy to reaccelerate the wings. This strategy effectively reduces the inertial necessary for flight [3, 4, 5, 6, 7, 8, 9]. Recent work directly measuring resonance properties in bees suggests that wingbeat frequencies are directly tuned to match resonance frequencies [10]. It remains an open question to what degree this resonance tuning is used among other insects and the potential implications of changing frequency. When wingbeat frequency is near the resonant frequency of the mechanical system, resonance may constrain wingbeats to a narrow range of energetically optimal frequencies [11, 12, 13]. With a resonance peak [14], small changes in wingbeat frequency may attenuate wingstroke amplitude. As a result, it has been proposed that insects are unlikely to modulate their wingbeat frequency on short timescales [15, 16]. On the other hand, if insects operate far from the resonance peak or if they require frequency control despite the potential power costs, then insects could freely modulate wingbeat frequency but would forego the energetic benefits of resonance. Because wingbeat frequency is an effective variable for adjusting flight forces, short time scale frequency modulation may play an underappreciated role in insect flight control, regardless of where insects operate on the resonance curve at steady state.

Insects have the capacity to change wingbeat frequency, regardless of whether they possess synchronous (neurogenic) or asynchronous (myogenic) flight strategies. In species with synchronous flight muscles such as *Manduca sexta*, wing motion is triggered by typically by a single action potential with sub-millisecond precision to the dorso-longitudinal muscles (DLMs) [17, 2, 18]. Although the nervous system can shift the timing of muscle activation [2], synchronous hawkmoths are often observed [4] and modeled [19, 20] as maintaining a constant wingbeat frequency. In contrast, asynchronous muscles are activated by mechanical stretch and pairs of asynchronous muscles antagonistically stretch each other to generate rhythmic wing motion [21, 22]. In this system, wingbeat frequency entrains to the resonance frequency of the wing-thorax system [23, 24]. To change wingbeat frequency, small accessory muscles must shift the resonance frequency of the thoracic structure [24], but this is a slow process because the accessory muscles only activate once per many wingstrokes [25]. For both synchronous and asynchronous insects, a dependence on resonance to reduce power requirements could limit the capacity for wingstroke-to-wingstroke changes in wingbeat frequency, which would require a deviation from the relatively slow changes in resonant frequency.

Despite the wingbeat frequency constraints imposed by resonance, modest changes in wingbeat frequency over long timescales are frequently observed in both asynchronous and synchronous insects. For asynchronous species, tethered *Drosophila melanogaster*, *Drosophila virilis*, and *Drosophila mimica* all exhibited a 10% change in wingbeat frequency in response to a slowly oscillating visual stimulus [11]. Under load lifting conditions, bumblebees (asynchronous) increased wingbeat frequency by approximately 5% between light and heavy load conditions, although an extreme individual changed frequency by 10% [26]. However, bumblebees do not change wingbeat frequency in 71 response to air density [27, 28]. In synchronous species, studies investigating wing damage [29], flight in turbulent flows [30], and varying forward velocities [31] in free flying *Manduca sexta* report 5-10% changes in wingbeat frequency. The results are consistent with 10% increase in wingbeat frequency when tethered locusts (synchronous) were flown in a wind tunnel at different wind speeds [32]. These studies suggest that modest changes in frequency may be a common method to adjust flight forces over the duration of many wingbeats.

Wingbeat frequency modulation at rapid timescales may be particularly useful for rejecting perturbations, but are especially challenging in a resonant system. To reject environmental perturbations, insects must adjust aerodynamic force production. Experiments in tethered *Drosophila* show that lift production increases with increased wingbeat frequency [11]. In the wingstroke-to-wingstroke regime, it remains unclear how insects balance the needs for economical and controllable flight. Therefore, insects could 1) forego wingbeat frequency modulation in favor of remaining on resonance to reduce power requirements or 2) trade off energy economy for increased control capacity in response to sudden perturbations. Given sharp resonance peaks in *Manduca sexta* [14], we hypothesize that wingbeat frequencies will remain constant even during maneuvers.

One of the major challenges in measuring wingbeat frequency modulation is in designing experiments to elicit behaviors that deviate from steady-state [33]. We were inspired by prior studies that used mechanical perturbations to trigger recovery maneuvers in free flying insects [34, 35]. In particular, prior work has shown that moths hover feeding from flowers can be successfully perturbed by vortex rings [35]. We adopted a similar experimental paradigm and tuned the strength of the vortex perturbation to be just weak enough for the moth to remain airborne. This setup enabled us to measure the steady-state behavior (hover feeding) to compare against the recovery maneuver, visualize the perturbations, and allowed the animals to freely maneuver.

Observing changes in wingbeat frequency does not necessarily alone reflect a change in the underlying oscillation frequency set by the nervous system of synchronous insects. Mechanical perturbations could physically move the wings to drive wingbeat frequency modulation without changing the underlying time-periodic forcing. We hypothesize that any observed modulation in wingbeat frequency are due to changes in underlying neural activation frequency. To address this hypothesis, we extracted instantaneous phase from wing kinematics and analyzed phase to assess changes in muscle input frequency [36]. In this framework, we used pre-perturbation kinematics to estimate phase if there was no perturbation. If this prediction differs from measured phase following a perturbation, we can conclude that an active change in motor input frequency must have occurred during the perturbation response.

## 2 Methods

**Animal care** *Manduca sexta* pupae raised on a diet containing retinoic acid were acquired from the University of Washington and Case Western Reserve University colonies. Adults were housed in an incubator on a 12:12 light:dark cycle. Each moth (n = 8) was used for one set of experiments ranging from 1 to 22 perturbations, depending on the willingness of the moth to return to feed from an artificial flower. Moths had no prior exposure to the artificial flower and were dark adapted for at least 30 min before experiments. Both males and females were used.

**Vortex ring perturbations** Perturbations (n = 58) were performed on adult *Manduca sexta* in a 1 cubic meter free flight chamber. The flower face was illuminated at 0.3 lux with a white LED, “moon” light (CW-126, Neewer, Shenzhen, China). Once the moth began feeding from the artificial flower, a lateral perturbation was generated via a vortex ring generator (MIGHTY Blaster Fog Ring Launcher, Incredible Science, USA) (Fig. 1 a). The vortex ring generator simultaneously produced artificial smoke composed of propylene gycol, glycerin, and distilled water (Super Zero Fog Fluid, Zero Toys, Concord, MA 01742) for vortex ring visualization. We saw no noticeable response in the moths to the approaching vortex ring. We were able to generate consistent vortex rings with an average diameter of (19.9 ± 0.6 cm; n = 10) with a velocity of (440 ± 30 cm s^−1^; n = 10). We positioned the vortex generate such that the impact was sufficient to knock most moths complete away from hover-feeding but not so great as to cause collision with the chamber walls. Moths were perturbed from the right posterior direction at constant elevation (Fig. 1 a) (Supplemental Movie S1).

**Figure 1.**
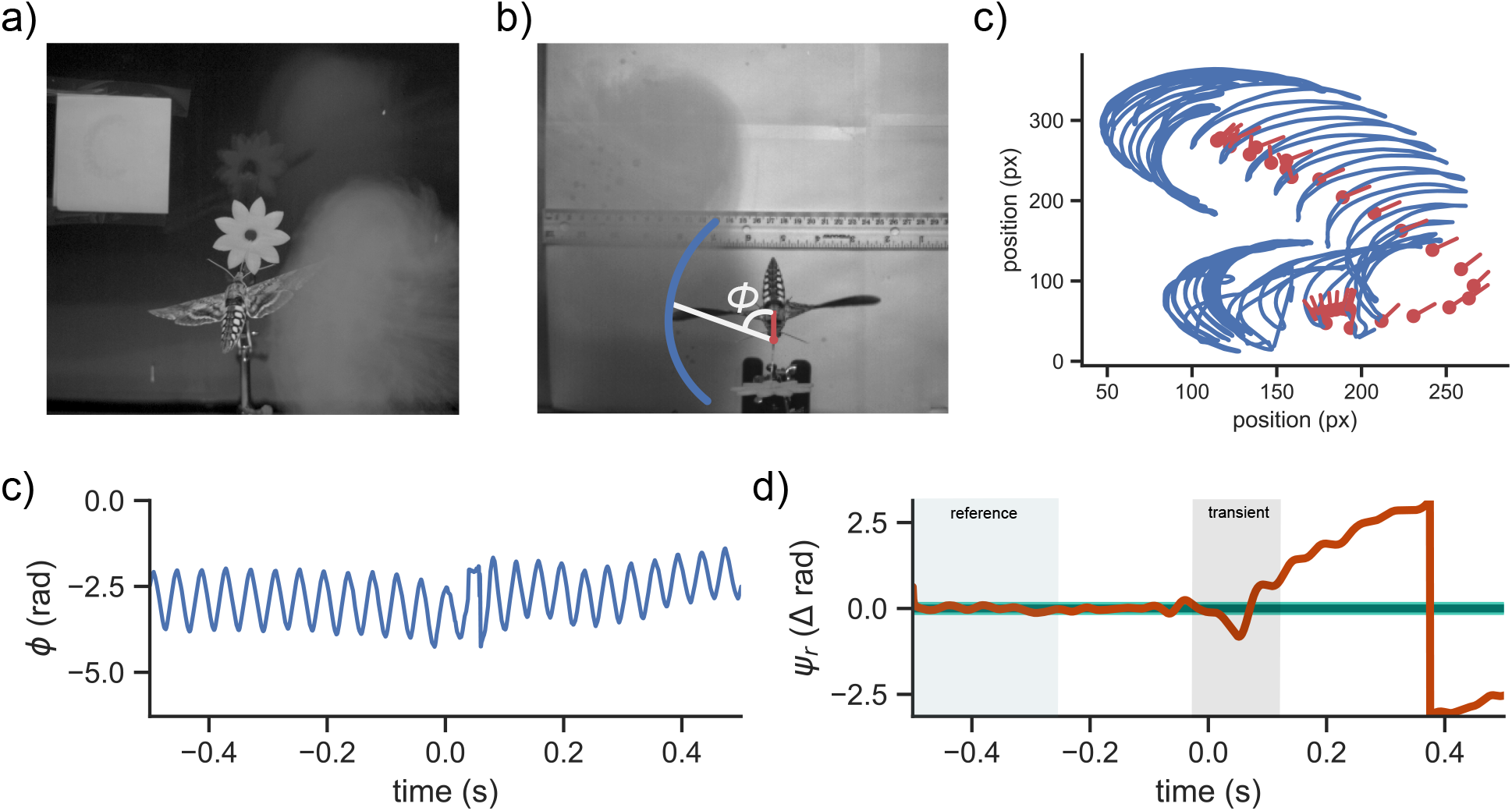
Vortex ring perturbations drive active recovery maneuvers in *Manduca sexta*. Representative analysis pipeline a) Rear view of a hawkmoth hovering in front of an artificial flower with an approaching vortex ring. b) Top-down view of a perturbation. Blue trace denotes approximate wingtip trajectory. Red dot marks proboscis base and red line points towards the thorax. *ϕ* is the instantaneous angle between wing position and body axis. c) Raw kinematic traces. Blue trace marks wing position. The red dot is the base of the proboscis and the red line points towards the center of the thorax. d) *ϕ* from the experiment in c). Perturbation occurs at t = 0s. d) Phase difference between actual phase and reference phase. Green rectangle highlights the region during which we build a reference phase and grey rectangle marks the approximate transient region of the perturbation. Light green region is the 95% confidence band of the reference phase while dark green is the mean projected phase. Orange line is measured phase. Perturbation occurs at 0s. In this example, we see a clear divergence in *ψ_r_* following the perturbation, indicating a Class 3 response.

**Kinematic extraction from high speed video recordings** All perturbations were filmed from above (dorsal view) at 2000 fps using a high speed camera (IL3, Fastec Imaging, San Diego, CA 92127). Infrared lights not visible to *Manduca sexta* provided illumination for high speed videography. We only analyzed videos where the tracked wing, head, and thorax remained in frame for ten wingstrokes post-perturbation.

To improve kinematic tracking fidelity, we background subtracted all video frames. We used DeepLabCut (ver. 2.1.1) to extract x-y position for the wingtips, thorax, and base of the proboscis (Fig. 1 b) [37] (Supplemental Movie S2). To train we used 7,111 frames split over 9 perturbations that were semi-manually digitized via DLTDdv [38] for 135,000 training iterations. Some perturbations had poor tracking performance because the moth flew over the flower or passed over the ruler. To refine the tracking, we used the refine labels function in DeepLabCut and manually digitized an additional 474 frames.

To quantitatively reject perturbations with poor tracking quality, we did not analyze any perturbation with over 1% of data points outside of the 95% CI of 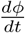. Qualitatively, this typically happens when a moth moves out of frame and the tracked points jump from one wing to the other.

### 2.1 Estimating wingbeat frequency from wing kinematics via instantaneous phase

Frequency modulation is a common perturbation rejection strategy, but it is difficult to quantify wingstroke-to-wingstroke changes in wingbeat frequency. We therefore adopt an instantaneous phase approach inspired by prior work on cockroach perturbation recovery [36].

To convert kinematic measurements to a continuous phase variable, we first defined a vector from the proboscis base to the thorax (Fig. 1 a). We defined a second vector from the thorax to the wingtip. We then calculated the angle (*ϕ*) between these two vectors (Fig. 1 c). We then applied the Hilbert transform (Eq. 1), which converts a real signal into the complex domain by convolving the signal with 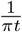.

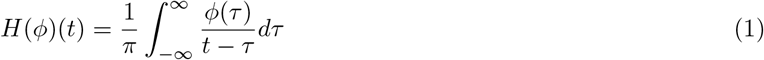

From here, we calculated the the phase variable *ψ*(*t*) as the complex angle of *H*(*ϕ*)(*t*). This method of estimating phase from time series measurements is well established in the biology literature [39, 40, 18].

Although calculating *dψ/dt* provides an estimate of instantaneous frequency, we still need to estimate wingbeat frequency from *ψ*(*t*). We attempted to ground our quantification in physiological constraints. The DLMs in *Manduca sexta* typically fire only once per wingbeat [41, 2, 18], suggesting that wingbeat frequency is driven by a single event. Therefore, we defined an arbitrary phase threshold and defined wingbeat frequency as the inverse of the time between threshold crossings. The period of any individual wingstroke may depend on our set threshold, but we find that overall magnitude of frequency modulation is robust to changes in phase threshold (Fig. S1).

### 2.2 Calculating wingbeat frequency modulation

To assess the magnitude of wingbeat frequency modulation, we calculated the width of the 95% quantile (97th percentile - 2.5th percentile) for each perturbation. Because we found that the first nine wingstrokes post-perturbation were statistically significantly different from the pre-perturbation mean, we divided each trial into three intervals: 1) wingstrokes −4 to −1, wingstrokes 1 to 4 and wingstrokes 5 to 8, where the perturbation occurs at wingstroke 0. Within each interval, we calculated the 95% quantile width for each perturbation. We then use the average 95% quantile width across perturbations as a measurement of wingbeat frequency modulation. We aligned each perturbation to wingstroke 0 by visually examining high speed videos to determine the frames at which moths were initially perturbed.

To assess whether there was significant individual-to-individual variation, we used a two-way ANOVA with individuals and intervals (pre-perturbation, post-perturbation wingstrokes 1-4, and post-perturbation wingstrokes 5-8) as factors. Normalized wingbeat frequency modulation (95% quantile width/mean wbf) was the dependent variable.

## 3 Results

### 3.1 Manduca sexta is capable of ± 16% wingbeat frequency modulation at the wingstroke-to-wingstroke timescale

In order to test wingbeat frequency modulation at the wingstroke-to-wingstroke timescale, we quantified wingbeat frequency modulation as mean width of the 95% quantile width of wingbeat frequency. Intuitively, this measurement quantifies the variation in wingbeat frequency within each perturbation. Consistent with prior literature, we find only a 1.28 ± 0.68 Hz 95% quantile width for the four wingstrokes immediately prior to perturbations [30]. This means that in steady state we would typically only observe a 1 Hz change above or below the mean flapping frequency.

In contrast, immediately after perturbations, moths must adjust aerodynamic force production to remain air-borne. Because hawkmoths may be operating on resonance, we hypothesize that any changes in aerodynamic force production are accomplished by methods other than wingbeat frequency modulation. Our data rejects this hypothesis. The 95% quantile width increased over 600% during the first four wingstrokes post-perturbation to 8.25 ± 3.96 Hz (*p* < 1E-18) (Fig. 2 c). This corresponds to a modulation range of 32%, which is comparable to previously reported magnitudes in other insects despite shorter timescales (Fig. 5 a). When we analyzed wingbeats 5-8 following the perturbation, we observed a decrease in the 95% quantile width to 2.88 ± 2.3 Hz (*p* = 0.03).

**Figure 2.**
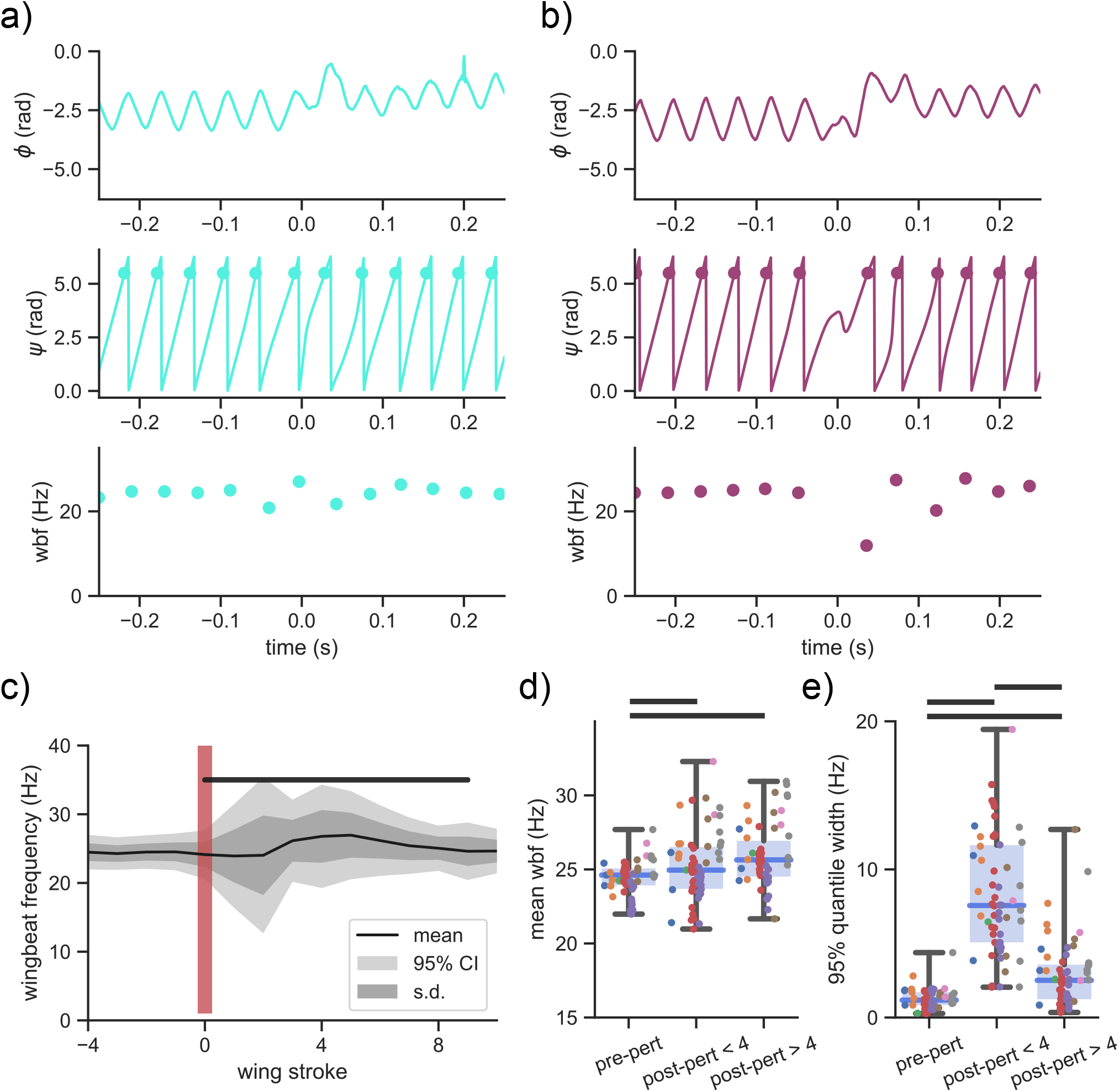
*Manduca sexta* exhibits rapid wingbeat frequency modulation immediately post-perturbation. a) representative analysis for a low frequency modulation perturbation. *ψ*, and wbf all remain constant despite the tracking eeror in *ϕ* at 0.2 s. b) representative analysis a lot frequency modulation perturbation. Fluctuations in *ϕ* propagate to *ψ* and wbf. c) mean, s.d., and 95% CI of wingbeat frequency for 58 perturbations. Red vertical line marks perturbation onset (wingstroke 0). Black horizontal bar denotes wing strokes statistically significant from pre-perturbation wingbeat frequencies. d) boxplots of mean wingbeat frequency for pre-perturbation (wing strokes −14 to −1), recovery response (wing strokes 1 to 4) and equilibrium response (wing strokes 5 to 8). Each color denotes an unique individual. e) 95% quantile mean wingbeat frequency for the same intervals as d). For d) and e), Black horizontal bars denotes statistical significance between intervals *p* < 0.05.

There was a large degree of variation across individual perturbation responses. Qualitatively we observed the largest modulation when the animals had the largest changes in their flight trajectory following perturbation. In a few perturbation responses, the degree of modulation did not exceed the steady-state variations (2 Hz at 5th percentile of perturbation responses). However, in the most extreme case, moths could change their wingbeat frequency by as much as 14.5 Hz (95th percentile of perturbation responses).

We also observed increases in mean wingbeat frequency following the perturbation. Before the perturbation, the mean wingbeat frequency was 24.4 ± 1.3 Hz. The first four wingstrokes post-perturbation increased to 25.2 ± 2.3 Hz (*p* = 0.01) and the subsequent four wingstrokes were also elevated at 25.9 ± 2.1 Hz (*p* < 1E-7). Normalizing wingbeat frequency modulation by these mean values results in modulation of ± 2.5%, ± 16%, and ± 5%, respectively.

In our two-factor ANOVA, we did find that individual moths had a weak effect (*p* < 0.01, *η* = 0.12) and individuals had different patterns of frequency modulation (interaction term; *p* < 0.01, *η*^2^ = 0.18), however independent of individual considerations, perturbations interval was still the largest determining factor for the frequency modulation (*p* < 1E-32, *η*^2^ = 0.63). By overlaying the raw datapoints for every perturbation and color coding individuals, our conclusion that *Manduca sexta* utilizes wingstroke-to-wingstroke wingbeat frequency modulation is supported by the data (Fig. 2 e).

### 3.2 Changes in wingbeat frequency reflect changes in motor activation frequency

The power of measuring instantaneous phase in a perturbation-based experimental paradigm is the ability to assess changes in neural drive frequency [42, 36]. Following the methods of Revzen et al. (2013), we first estimated a reference clock frequency and projected this oscillator forward in time by fitting a regression line to *ψ* from −0.5 s to −0.25 s. From here, we subtract the measured phase from the reference phase to calculate the phase residual (*ψ_r_*) (Figs. 3 & 1 c). Residual phase shows how much the perturbation has caused the animal to deviate from the phase it would have had if there was no perturbation. Following a perturbation, there are three classes of possible *ψ_r_* outcomes: 1) no difference (Fig. 3 a), 2) constant phase offset (Fig. 3 b), and 3) continuous phase divergence (Fig. 3 c).

**Figure 3.**
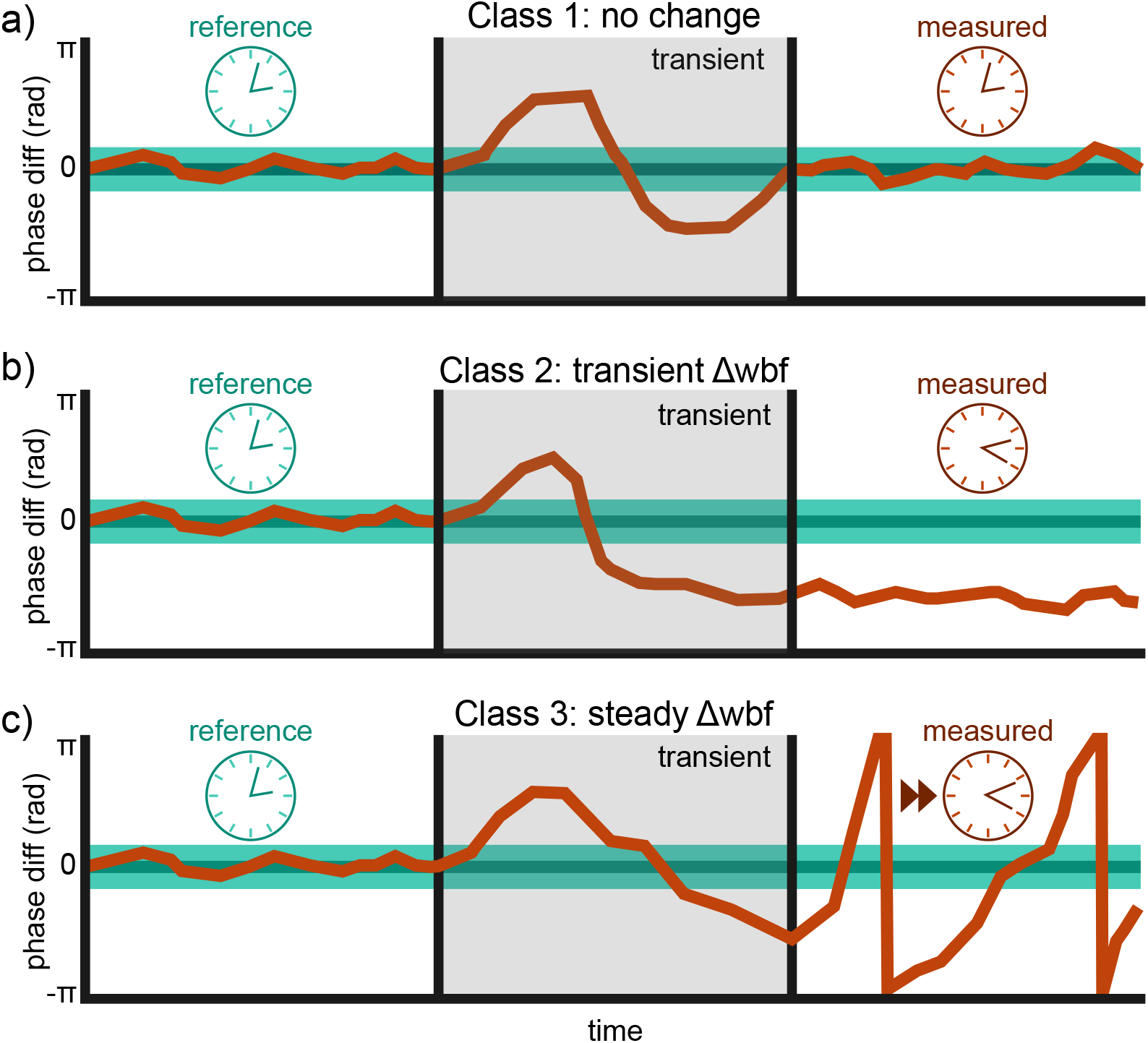
Schematic of instantaneous phase analysis and possible responses. Comparing post-perturbation to a pre-perturbation reference phase enables quantification of neural changes. a) Class 1 response where *ψ_r_*(*t*) returns to the pre-perturbation phase. Does not require a change in motor input because the underlying clock frequency remains constant. b) Class 2 response where *ψ_r_*(*t*) = *c* requires a transient change in motor input. The underlying clock needs to change frequency to achieve a constant phase offset. c) Class 3 response where *ψ_r_*(*t*) continuously diverges requires a lasting change in motor input. For all panels, dark green line denotes projected reference phase, while light green bands are the 95% CI of the projection. Orange line is the hypothesized measured phase.

It is possible that any transient frequency modulation is a purely passive response to the perturbation and not necessarily due to a changes in muscle activation frequency [36]. To address this uncertainty, we utilized instantaneous phase analysis to quantitatively assess changes in neural input frequency. We divided perturbations into three response classes. Class 1 responses exhibited no lasting change in *ψ_r_*(*t*). This would indicate that underlying frequency modulation was not necessary and is consistent with the insects responding purely mechanically and pacing the flight muscles at a constant frequency (Fig. 4 a). Class 2 responses are characterized by a constant *ψ_r_*(*t*) offset after the initial transient (Fig. 4 b). To achieve a lasting phase difference in an oscillatory system, there must be at least a temporary increase or decrease in phase velocity (i.e. frequency). Therefore, a Class 2 response requires a transient wingbeat frequency modulation. Finally, Class 3 responses have a continuously diverging *ψ_r_*(*t*). Diverging phase requires that the moth adopted a new pacing frequency in response to the perturbation (Fig. 4 c). Importantly for our hypothesis, both Class 2 and Class 3 responses require active neural change in wing beat frequency.

**Figure 4.**
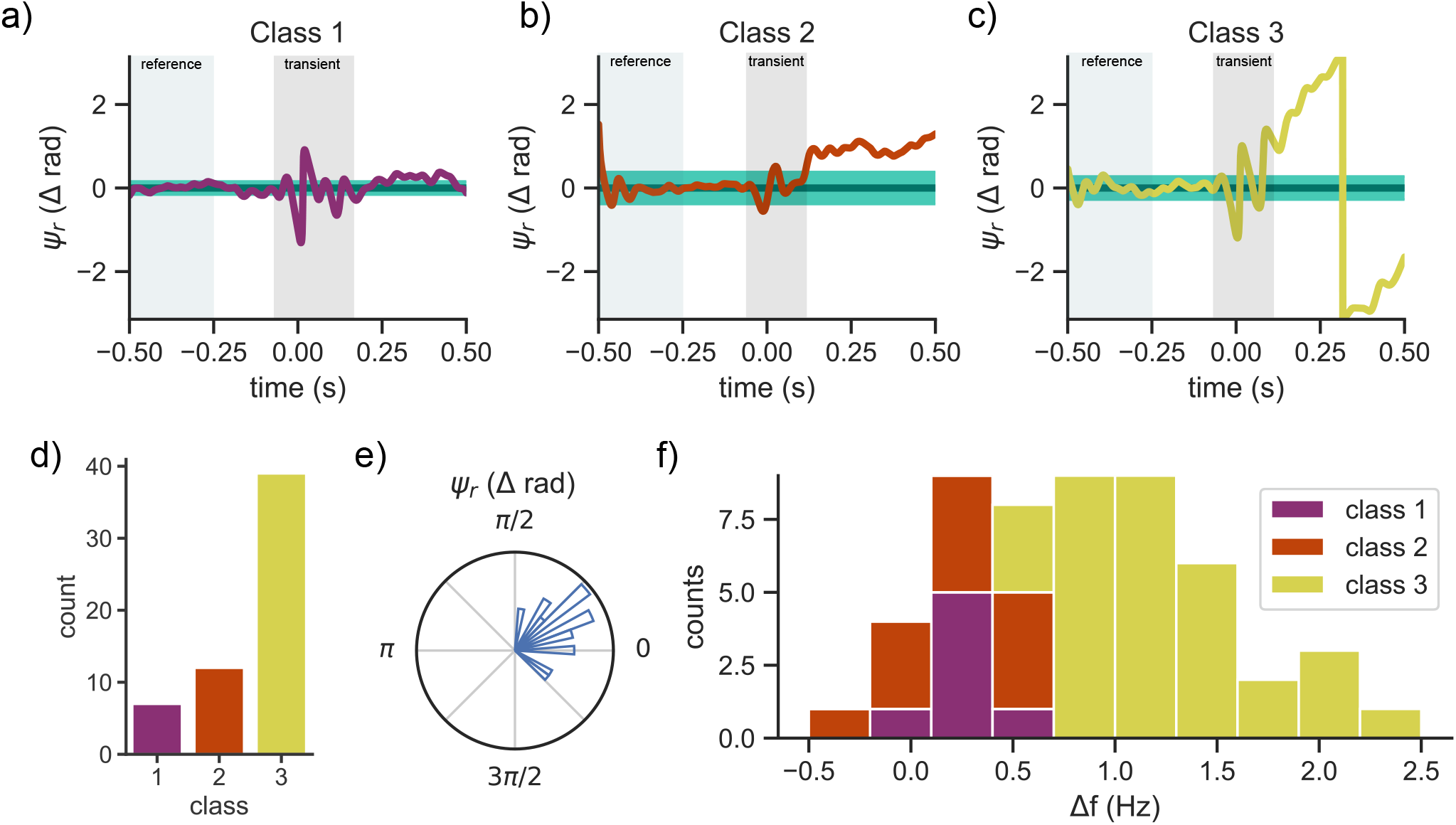
Over 85% of perturbations require a change in neural input frequency. Representative *ψ_r_*(*t*) for a) Class 1, b) Class 2, and c) Class 3 responses, where the perturbation began at 0 s. Green horizontal bar denotes the mean and 95% CI of the projected phase. Purple, orange and yellow traces are measured *ψ_r_*(*t*). Light green and grey rectangles mark the regions where we build the reference phase and the transient perturbation response. d) Counts of class responses. e) Circular histogram of *ψ_r_*(*t*) for Class 1 and Class 2 responses. Density is denoted by area instead of radii. f) Histogram of mean frequency change for all three classes calculated as the slope in *ψ_r_*(*t*) slope from 0 to 0.5 s. This Δ*f* represents mean changes in wingbeat frequency over 0.5 s and not wingbeat-to-wingbeat frequency modulation.

**Figure 5.**
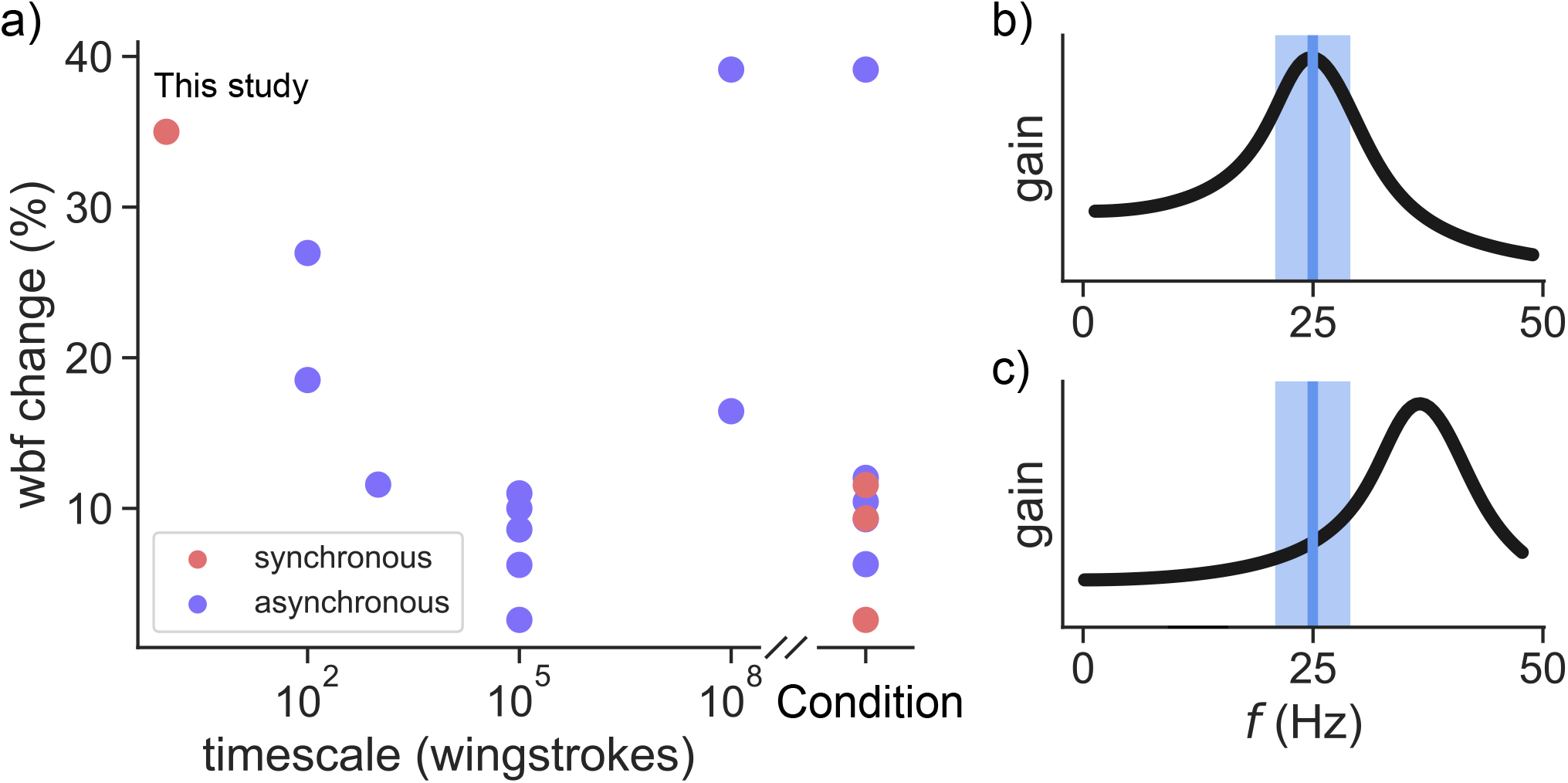
a) brief survey of literature comparing wingbeat frequency modulation and the timescales of these changes. “Condition” refers to studies where wingbeat frequency change is observed over discrete conditions an indeterminate experimental intervals. The synchronous leftmost point is from data collected from *Manduca sexta* in this manuscript. Other data was aggregated from [29, 30, 31, 8, 50, 62, 27, 51, 26, 52, 53, 11, 54]. b) and c) possible resonance curves with mean wingbeat frequency (blue vertical line) and modulation (blue rectangle). b) resonance matches typical wingbeat frequencies. c) resonance frequency is 50% greater than typical wingbeat frequencies.

To distinguish between classes, we first fit a line (*y* = *mx* + *b*) to *ψ_r_*(*t*) from 0 to 0.5 s following the perturbation (accounting for the wrapping of phase at *π* and −*π*). Prior literature has found a roughly 5% change in wingbeat frequency in *Manduca sexta* over longer timescales [30, 31, 29]. Therefore, we set a frequency change threshold of ±2.5% (±0.625 Hz). To distinguish between Class 1 and Class 2 from Class 3 responses, we tested if there was a significant slope to the residual phase line after the perturbation. To distinguish Class 1 and Class 2 responses, we determined if there was a significant change in the residual phase. To do so, we looked at the average value of the line (i.e. at 0.25 s). If this value was outside of the 95% confidence interval of our projected phase, we conclude that a phase shift must have occurred (Class 2 response). Otherwise if there is no significant slope and no significant phase residual, we classify the response as Class 1. We used the following criteria to distinguish between classes:

1. Class 1: |*m*| < 0.625 Hz, 5% CI < b < 95% CI
2. Class 2: |*m*| < 0.625 Hz
3. Class 3: |*m*| > 0.625 Hz

Of the 58 perturbations we recorded, over 85% exhibited a Class 2 or 3 responses (Fig. 4 d), which all require a change in the frequency of the underlying neural commands. Regardless of where changes in motor commands arise in the nervous system, we conclude that changes in observed wingbeat frequency are driven by active modulation of motor input.

For Class 1 and Class 2 responses, it is possible that *Manduca sexta* randomly adopts a new phase. Because we do not control for the timing of when the perturbation impacts the wing, a uniform distribution of post-perturbation phase would indicate that timing has no effect on the recovery response. We would also be able to test whether Class 1 responses are a subset of Class 2 responses that adopted the same initial phase. To test this hypothesis, we analyzed the post-perturbation phase for the aggregate of Class 1 and Class 2 responses. Using a Rayleigh test [43], we found that post-perturbation phases were not uniformly distributed (*p* < 1E-6; Fig. 4 d)). Instead, we report a vector strength of 0.87 with a mean phase of 0.49 rad, where a vector strength of 1.0 is perfect phase synchrony (Fig. 4). Therefore perturbation timing is insignificant and Class 1 responses are distinct from Class 2 response. The preferred phase post-pertubation may reflect specific patterns of frequency modulation underlying the recovery to vortex ring perturbations. The simplest explanation is the persistent phase lead comes from a transient increase in wingbeat frequency – the moths take a few faster wingstrokes during perturbation recovery. This would be consistent with the need to create overall more aerodynamic force to accomplish perturbation recovery than steady hovering. However, a phase advance alone could also result from more complex patterns of frequency modulation.

## 4 Discussion

### 4.1 *Manduca sexta* utilizes substantial wingbeat frequency modulation during maneuvers

Given the presence of a resonance peak [14] and observations of nearly constant wingbeat frequency in *Manduca* flight [4], we had hypothesized that hawkmoths would abdicate wingstroke-to-wingstroke frequency modulation in exchange for reduced power requirements. Instead, our data rejected this hypothesis and showed that *Manduca sexta* can adjust wingbeat frequency by an average of 32% at the wingstroke time scale. The capacity of some moths was even greater. One measure of frequency modulation capacity is to take the most extreme example of each individual, which corresponds to a 12.7 Hz bandwidth or 50% modulation range. In this case *Manduca sexta* has the capability to range from ~ 19 to ~ 31 Hz within four wingstrokes.

In support of our second hypothesis, we found that over 85% of perturbations required active changes in motor input frequency. Wingbeat frequency modulation over long timescales is a common strategy to respond to changing aerodynamic requirements [44, 11, 30]. Extending on this work, our results indicate that wingbeat frequency modulation is an active control strategy for *Manduca sexta* at the wingstroke-to-wingstroke timescale (Fig. 5 a). This approach was used previously to show that cockroaches respond to large lateral perturbations first with an initial 100 ms that is consistent with just mechanics (no frequency change necessary). This initial response is followed by a longer, neurally mediated reaction during which there is a consistent 5% decrease in stride frequency [36].

Given the oscillatory nature of flapping wing flight, changes in wingbeat frequency have significant implications for power requirements. Because inertial and aerodynamic power requirements both scale with frequency cubed [24], increasing wbf by 16% would increase mechanical power requirements by over 55%. Spring-like structures in the insect flight system have a finite capacity for elastic energy exchange [6], therefore the musculature would have to supply this increased power demand. Under tethered *in vivo* conditions, *Manduca sexta* power muscles produce only 50% of peak power and this power can be modulated with precise timing [41, 2]. These experiments suggest that the muscles have excess capacity to drive elevated wingbeat frequencies.

### 4.2 Wingstroke-to-wingstroke frequency modulation complements a broad suite of control strategies in *Manduca sexta*

In *Manduca sexta*, there are many mechanisms for overcoming perturbations. By virtue of having flapping wings, *Manudca sexta* can dampen rotational perturbations with no active change in kinematics via a concept called flapping wing counter torque (FCT) [45]. The premise of FCT is that any body rotation increases drag in the opposite direction because the wings are continuously oscillating. Notably, both active aerodynamic force production and flapping wing counter torque increase with wingbeat frequency. Therefore, increasing wingbeat frequency simultaneously increases maneuverability and stability, or vice versa [45].

In addition to passive control mechanisms, wingbeat frequency modulation must work in tandem with active changes in muscle activation. In *Manduca sexta*, wing kinematics are controlled by a set of 10 muscles, so any changes in wingbeat frequency likely affects the coordination of these muscles. In *Manduca sexta*, the downstroke is triggered by typically a single action potential to the left and right DLMs [46, 2, 18]. Precise changes in the activation phase of these DLMs can account for 50% of yaw torque production [2]. Because of the strong phase dependence of *Manduca sexta* DLMs [2], the nervous system may need to maintain precise control over muscle activation phase while the frequency of muscle strain changes from wingstroke-to-wingstroke. Further complications arise because recent work has shown that the timing of the 10 major muscles controlling wing kinematics are tightly coordinated in their activation timing [18]. Changes in wingbeat frequency may require changes in activation frequency for all ten muscles involved orchestrated through phase shifts across the entire motor program.

Insects can respond to perturbations at the wingstroke timescale [47, 48, 49, 34, 40], but frequency changes typically happen over many wingstrokes (Fig. 5 a) [29, 30, 31, 8, 50, 27, 51, 26, 52, 53, 11, 54]. The relatively slow wingbeat frequency changes in asynchronous insects may be because wingbeat frequencies entrain to the resonant frequency of the wing-thorax system. Changes to resonance frequency are thought to be driven by the pleurosternum muscle [24], which stiffens the thorax, but only activates once per many wingstrokes [25]. Our study bridges the gap between these two bodies of literature by showing that *Manduca sexta* utilizes wingbeat frequency modulation at the wingstroke timescale to recover from transient perturbations.

### 4.3 Resonant systems cannot escape tradeoffs between energy and control

Many authors have stated that insects are tuned to resonance [15, 24, 14, 22, 16, 10] (Fig. 5 b) due to observations of constant wingbeat frequencies and estimates [14] and direct measurements [10] of a dominant resonance peak. This is consistent with evidence that elastic energy exchange substantially reduces the power requirements of insect flight [3, 4, 5, 6, 7, 8]. As the wing decelerates, elastic elements store excess kinetic energy and subsequently return this energy to reaccelerate the wing. Critically, this elastic energy exchange process has an intrinsic timescale (resonant frequency). Therefore, efficient energy exchange requires an insect’s wingbeat frequency to match the resonance frequency of the wing-thorax system. Even in the case where wingbeat frequencies are off resonance, large amplitude changes in wingbeat frequency can have significant consequences in a resonant system.

A dominant resonant peak introduces frequency dependent mechanics (Fig. 5 b & c). If steady-state wingbeat frequencies are tuned to resonance [A dominant resonant peak introduces [15, 24, 14, 22, 16, 10], elastic recoil of the wing-thorax system would increase the ratio (i.e. gain) between output wing amplitude to input force (Fig. 5 b). Critically, this gain is itself frequency dependent so small deviations in wingbeat frequency could cause large changes in wingbeat amplitude [14]. Both increases and decreases in wingbeat frequency would lead to reduced wingbeat amplitude unless the muscles generated additional power. Consistent with this perspective, metabolic rate increases with increased wingbeat frequency in bumblebees [26]. On the other hand, operating away from resonance could lead to a monotonic relationship between frequency and gain (Fig. 5 c). This could enable frequency modulation as a knob to control wingbeat amplitude (Fig. 5 c). Indeed, both wingbeat frequency and amplitude are correlated with lift production in flies [8]. We observed a 32% range in wingbeat frequency in *Manduca sexta*, which suggests that resonance either 1) adds potential energetic costs when wingbeat frequencies deviate during perturbation recovery (Fig. 5 b) or 2) amplifies the control potential of wingbeat frequency modulation (Fig. 5 c). These hypothesized resonance curves provide a qualitative view of the potential tradeoffs between energy and control faced by hawkmoths trying to use resonant, flapping wing flight.

The qualitative resonance implications in insects have been quantified in their robotic counterparts. In insect-scale flapping wing vehicles, tuning wingbeat frequencies to resonance improved energy efficiency by 50% [9]. However, the energetic benefits of resonance also restricted the range of attainable wingbeat frequencies to just ± 5% around resonance [13]. Due to these constraints, controlled flight was achieved by modulating stroke amplitude but explicitly holding wingbeat frequency constant [55]. In addition, resonance can filter out subtle wing kinematics that are important in producing aerodynamic forces [13, 8, 11, 56, 57]. By moving off resonance, yaw rate improved 10-fold, but required a 40% increase in voltage [13]. These results in robotic systems quantify the extreme tradeoffs between energetic cost and control in insect-scale flapping wing systems.

Beyond flapping wing flight, combining actuation and elasticity poses significant dual challenges for control and energetics in other forms of locomotion. For example, elastic elements in swimming cetaceans [58] and running humans [59] introduce frequency dependent energetic costs. Although energetically beneficial, changes in frequency require changes to elasticity [59] or center of mass [60]. Extending to impulsive movements, springs are necessary to overcome constraints of power limited muscles [61]. However, control over these power amplified movements is limited to careful design of latch mechanisms because once the energy is released from storage, control would require a great deal of rapid power production. A unifying challenge for impulsive and oscillatory systems is that potential energy can be beneficial for certain performance measures, but control becomes challenging when there is too much potential energy.

